# *CHRFAM7A* overexpression in human iPSC-derived Interneurons dysregulates α7- nAChR surface expression and alters response to oligomeric β-amyloid peptide

**DOI:** 10.1101/2024.06.04.597325

**Authors:** Maria Llach Pou, Camille Thiberge, Stéphanie Pons, Uwe Maskos, Isabelle Cloëz-Tayarani

## Abstract

The α7 neuronal nicotinic receptor (α7-nAChR) gene, *CHRNA7*, is widely expressed within the brain and at the periphery. It plays various important roles in cognition and immune functions. Decreased expression of α7-nAChR has been associated with Alzheimer’s disease (AD) triggered by the accumulation of the 42-amino acid beta-amyloid peptide (Aβ_1-42_). The interactions of this peptide with α7-nAChR may represent a pivotal mechanism that is involved in pathogenesis of AD. The regulation of *CHRNA7* is a complex process. Normal function of α7-nAChR in mammalian cells requires the co-expression of chaperone proteins such as RIC3 and NACHO which facilitate the formation of cell surface receptors. In humans, *CHRNA7* regulation also involves the specific chimeric *CHRFAM7A* gene product dupα7, which may assemble with α7 subunits and lead to dominant negative regulation of α7-nAChR function. To further elucidate the complex interplay between *CHRFAM7A* gene product (dupα7), α7-nAChRs and Aβ_1-42_, we used human induced pluripotent stem cells (iPSC)-derived interneurons (INs). Four iPSC lines were analyzed for the presence of *CHRFAM7A* copies. Among them, a cell line with a null genotype was selected for the lentiviral overexpression of *CHRFAM7A*. Our data show that overexpression of *CHRFAM7A* led to a reduction in the surface detection of α7-nAChR ligand binding sites in iPSC-derived INs. INs expressing the α7-dupα7 subunit (α7-dupα7-INs) exhibited lower levels of RIC3 and NACHO. Upon agonist treatment by nicotine, an up-regulation of α7-nAChR ligand binding sites was observed in α7-dupα7-INs as compared to non-transduced INs (α7-INs). At low levels of Aβ treatment, α7-INs displayed a significant reduction in production of reactive oxygen species (ROS), while high levels resulted in a slight increase. In contrast, α7-dupα7-INs exhibited lower baseline levels of ROS that remained unaltered by Aβ treatment. ROS are known to exacerbate AD pathogenesis. We hypothesize that such effects may also be triggered by α7-dupα7-INs in the brain of patients. Further investigations are currently undertaken to confirm this hypothesis.

## 1/ Introduction

The α7 nicotinic acetylcholine receptor (α7-nAChR) belongs to the family of pentameric ligand-gated ion channels. This receptor presents a rapid desensitization and high permeability to Ca^2+^, it is involved in neuronal plasticity, survival processes, and it activates a large number of signaling cascades thereby promoting synaptic transmission (Borroni & Barrantes, 2021). In the mammalian brain, the α7-nAChR is predominantly expressed in pivotal brain regions such as the cerebral cortex and hippocampus, which underlie the cognitive functions and memory processes (Borroni & Barrantes, 2021; Gotti et al., 2006; Potasiewicz et al., 2021). The α7-nAChR has been shown to contribute particularly to the plasticity of neuronal networks as well as to synaptic plasticity in the prelimbic cortex (Udakis et al., 2016). Other studies have shown that α7-nAChR in hippocampus is predominantly expressed by GABAergic inhibitory neurons and can regulate their activities through their presynaptic modulation of GABA release (Pidoplichko et al., 2013).

In addition, α7-nAChR has been shown to regulate adult neurogenesis and neural stem cell differentiation that occurs more specifically in the dentate gyrus of the hippocampus, and it also participates in reorganization of neuronal circuits upon reintegration of the newborn neurons (Catavero et al., 2018; Nacer et al., 2021; Rosato-Siri et al., 2006). Cerebral cortex and hippocampus are two brain structures that are involved in the pathogenesis of Alzheimer’s disease (AD) (Fjell et al., 2014; Geula, 1998). These structures express high densities of α7-nAChR which are therefore expected to participate in the molecular and cellular mechanisms involved in pathogenesis of this disease. Although this hypothesis has been confirmed by a large number of studies, the precise role of α7-nAChR in pathogenesis of AD is still unclear. Amyloid β peptide (Aβ) is considered to be one of the principal factors that triggers AD, via its accumulation, aggregation, and lack of clearance. Aβ is the primary component of senile plaques in AD patients also found in soluble form. Because of its initial accumulation within the cell, Aβ is thought to be responsible for the onset of AD (Ma & Qian, 2019; Volloch & Rits-Volloch, 2023). Additional studies have further explored the link between Aβ and α7-nAChR. It has been reported that reduced expression of α7-nAChR correlates with Aβ accumulation and with the associated cognitive impairments (Ma & Qian, 2019; Schliebs & Arendt, 2011). Other authors have shown that Aβ increases the trafficking of α7-nAChR to the plasma membrane (Liu et al., 2013) and that intracellular accumulation of Aβ in neurons is facilitated by α7-nAChR in AD (Nagele et al., 2002). The interaction of Aβ with α7-nAChR has proven to be a complex question. At physiological concentrations (pM to low nM range), Aβ triggers the desensitization of α7-nAChR in a conformation which is still sensitive to agonists. Whereas at higher concentrations (nM to low μM range), Aβ acts as a negative modulator resulting in opposite signaling pathways, neuroprotective vs. neurotoxic (Di Lascio et al., 2022; Lasala et al., 2019). In the human brain as compared to experimental cellular models which have been used so far, one can expect a more complex condition given the fact that the regulation of the gene *CHRNA7* which encodes α7-nAChR is not yet fully understood (Sinkus et al., 2015). In the human genome, the α7 subunit is encoded by *CHRNA7* gene which is located on chromosome 15 and comprises 10 exons (Gault et al., 1998). The human *CHRFAM7A* gene results from a partial duplication of exons 5 to 10 of the parent *CHRNA7* gene (Sinkus et al., 2015). *CHRFAM7A* is not found in rodents and non-human primates (Sinkus et al., 2015). *CHRFAM7A* was first studied in cell lines and heterologous systems and shown to translate into the so-called dupα7 protein (Wang et al., 2014). Dupα7 can assemble and form heteromeric α7-nAChR/Dupα7 with altered pharmacological properties as compared to α7-nAChR (Wang et al., 2014). When incorporated into the receptor in-vitro and in-vivo, data have shown that dupα7 downregulates the activity of α7-nAChR mainly due to a reduction of the number of ACh binding sites (Araud et al., 2011; Lucas-Cerrillo et al., 2011). Finally, the *CHRFAM7A* gene exhibits copy number variation (CNV) in the human genome (Szigeti et al., 2020). Its regulatory role on the α7-nAChR needs to be further investigated using appropriate experimental models close to human brain, taking into consideration that the general population presents variable numbers of *CHRFAM7A* alleles.

The objective of our study was to further determine the role of dupα7, the *CHRFAM7A* gene product, and the consequences of its interaction with α7-nAChR using human induced pluripotent stem cell (hiPSC)-derived interneurons (INs). We first developed a protocol to derive the hiPSC into MGE-derived interneurons. We analyzed the effect of dupα7 on some of the mechanisms involved in α7-nAChR trafficking to the neuronal membrane, and their reactivity to Aβ_1-42_. We first showed that dupα7 overexpression modifies the pharmacological properties of α7-nAChR expressed by human INs as well as their response to the toxic effects of Aβ_1-42_. We also evidenced some protective effects of dupα7 against oxidative damage.

## 2/ Experimental procedures

### 2.1. Culture of iPSC

Human iPSC cells (WTSli002-A, Wellcome trust) were maintained on Vitronectin XF™ (07180, Stemcell Technologies) coated 6-well plates (ref. 27147, Stemcell Technologies) with mTesR plus medium (ref.100-0276, Stemcell Technologies) at 37ºC, 5% CO_2_ and 5% O_2_. When 60-70% confluency was reached, cells were passaged as aggregates using 0.5 mM EDTA for 3 minutes at 37ºC. Absence of genomic abnormalities was assessed with the iSC-digital™ PSC test (Stemgenomics).

### 2.2. Lentiviral construction, transduction and cloning

To create the pLV-PGK-CHRFAM7A-IRES-tdTomato-WPRE vector, we replaced the sequence coding the β2 nicotinic subunit integrated into the pLV-PGK-β2-IRES-TdTomato-WPRE vector described previously (Koukouli et al., 2016) by the *CHRFAM7A* sequence taken from the plasmid pLVx-CHRFAM7A-IRES-ZSGreen (a kind gift from Dr A Baird from the Department of Surgery, University of California). Briefly, the fragment containing the *CHRFAM7A* sequence and part of the IRES sequence were excised with the restriction enzymes XhoI and AvrII. This fragment was then cloned using the same restriction sites in the pLV-PGK-β2-IRES-TdTomato-WPRE after excision of the ß2 subunit.

For Lentiviral transduction, 60-70% confluent iPSCs were infected with pGK-CHRFAM7A- ires-TdTomato lentivirus (MOI 15, 24h incubation). After three passages, labeled iPSCs were purified by FACS sorting and plated in 96-well plates (ref.38044, StemCell Technologies) at a density of 1 cell per well. Wells with more than one cell were discarded. Clones were firstly amplified in the 96-well plates and passaged as aggregates to gradually bigger surfaces.

### 2.3. Interneuron differentiation

After thawing, iPSCs were passaged at least three times prior to differentiation. The iPSCs colonies were firstly detached from the plate with 0.5 mM EDTA (3 min, 37°C) and broken manually by gently pipetting several times to obtain a homogeneous suspension. About 600,000 cells were seeded in one well of AggreWell™ 400 24-well plate (ref.34411, Stemcell) pre-treated with Anti-Adherence Rinsing Solution (ref.07010, Stemcell). To increase cell survival, 10 μM ROCK inhibitor (ref.72304, Stemcell Technologies) was added to the iPSCs cultures 30 minutes before treatment with 0.5 mM EDTA, in the medium for the first 24h of embryoid body formation (EB). During the first two weeks of differentiation, floating EBs were grown in SRM medium (15% Knockout serum replacement (ref.11520366, Thermo Fisher), DMEM (ref.11965092, Thermo Fisher), and 10 μg/mL Penicilin-Streptomicin) supplemented with 2 mM L-Glutamine (ref.07100, Stemcell) and 10 mM 2-Mercaptoethanol (ref.11528926, Thermo Fisher). The medium was partially changed every other day extremely slowly to prevent disturbing the EB formation. As described in Ni et al. (2019), for induction of neuroectoderm, EBs were treated with 10 μM SB431542 (ref.72234, Stemcell Technologies) from day 0 to day 7 and with 100 nM LDN193189 (ref.72147, Stemcell Technologies) from day 7 to day 14. For the induction of an MGE phenotype, cells were treated with 100 nM SAG (73414, Stemcell Technologies) from day 0 to day 21, 5 μM IWP2 (ref.72124, Stemcell Technologies) from day 0 to 7 and 50 ng/mL FGF8 (ref.100-32, Peprotech) from day 14 to day 21. At day 14, EBs were filtered (ref.27215, StemCell Technologies), seeded in PLO/Lm coated 6-well plates (EBs produced in one well of AggreWell™ 400 24-well plate were usually seeded into one well of a 6-well plate) and maintained in AAN2 medium (0.5% N2 Supplement (ref.07152, Stemcell Technologies), 200 mM AA-ascorbic acid (ref.12084947, Thermo Fisher) and 10 μg/mL Penicilin-Streptomicin). At day 21, cells were dissociated with Trypsin-EDTA 0.05% supplemented with 100 mM Trehalose (5 min, 37°C) and any remaining EB aggregate was dissociated manually by pipetting. An equal volume of DMEM supplemented with 10% FBS and 2 U/mL Turbo DNAse was added to inhibit trypsin and incubated for 15 minutes to remove any sticky DNA mass released. Cells were then centrifuged, resuspended in freezing media (10% DMSO, 100 mM Trehalose, FBS) and split in cryovials containing 4-5 million cells. Cells were harvested slowly overnight in a Nalgene® Mr. Frosty box at -80ºC and transferred to a liquid nitrogen tank after 24h. One week before the experiment, cells were thawed quickly at 37ºC and seeded in PLO/Lm coated surfaces at a density of 100.000 cells / cm^2^. The cells were maintained in B27GB medium (1% B27 Supplement (ref.11500446, Thermo Fisher), DMEM-F12 (ref.1554546, Thermo Fisher) and 10 μg/mL Penicilin-Streptomicin). From day 21 to day 28, B27GB medium was supplemented with 5 ng/mL GDNF (78058, Stemcell Technologies) and 5 ng/mL BDNF (ref. 78005.1, Stemcell Technologies) and 1% CultureOne Supplement (15674028, Thermo Fisher). In addition, cells were treated with 2 μM PD0332991 (ref. PZ0199-5MG Sigma Aldrich) and 2.5 μM DAPT (D5942-5MG, Sigma Aldrich) to induce postmitotic phenotype.

### 2.4. Nicotine treatment

Nicotine (ref N3876, Sigma) was diluted in phosphate buffer (pH 7,4) to achieve a concentration of 100 mM. To treat the cultures, 1 mM Nicotine diluted in B27GB medium (50% fresh + 50% conditioned) was used. IN progenitors were exposed to nicotine for 30 min (acute treatment) or 3 days (chronic treatment).

### 2.5. Aβ_1-42_ treatment

Aβ_1-42_ concentrated stocks were prepared prior to the experiments. Briefly, 1 mg of HFIP-treated Aβ_1-42_ (ref. 4090148.0500, Bachem) was resuspended in 100 μL NH4OH 1% and 900 μL Tris/NaCl buffer (50 mM Tris pH7,6 NaCl 250 mM) to achieve a 220 μM concentration. The resuspended Aβ_1-42_ peptide was aliquoted, flash frozen (in liquid nitrogen) and stored at -80°C until use. The day of the experiment, Aβ_1-42_ peptide was thawed on ice and serial dilutions were done to achieve 1000 nM, 100 nM, and 0.1 nM concentrations in B27GB medium (100% conditioned). The Aβ_1-42_ peptide was added to the cultures and incubated for 2h at 37°C.

### 2.6. Quantitative Real-Time PCR (qRT-PCR)

For gene expression studies, the total mRNA from neuronal cultures was extracted with the RNeasy Plus Mini Kit (ref. 74034, Qiagen). cDNA was generated using the High-Capacity cDNA Retro-Transcription Kit, according to the manufacturer’s instructions (ref. 4368814, ThermoFisher). qRT-PCR was performed on the Applied Biosystems 7500 Real-Time PCR System. UDG activation (1 cycle, 2 min, 50°C) and polymerase activation (1 cycle, 10 min, 94°C) were followed by 40 cycles of denaturation (15s, 95°C) and annealing / extension (1 min, 60°C). The PCR reactions contained 500 nM of each primer, 0.5 × PowerUP SYBR Green Master Mix (2X) (ref. A25742, Applied Biosystems), and 10 ng of cDNA in a final volume of 10 μL. Primer sequences are presented in Suppl. Table S1. The specificity of the primers was verified with a melting curve and agarose gel electrophoresis of the PCR content confirmed amplicon size. The mRNA expression level was expressed as the fold change to the housekeeping gene GAPDH using the 2−ΔCT method.

### 2.7. CNV calculation

For copy number variation (CNV) analysis, gDNA was extracted from neuronal cultures with the GenEluteTM Mammalian Genomic DNA Miniprep Kit (G1N10, Merck). qPCR was performed as described above with 20 ng of gDNA in a final volume of 10 μL. For each pair of primers, two standard curves with Log10 gDNA dilutions were performed to determine amplification efficiency. Primer sequences are presented in Suppl. Table S2. The control gene RNAseP and a control cell line (a kind gift from Dr E Bacchelli, FaBiT – Department of Pharmacy and Biotechnology, Bologna, Italy) were used to calculate CNV using the Pfaffl method (see Suppl. Fig.1).

**Figure 1:**
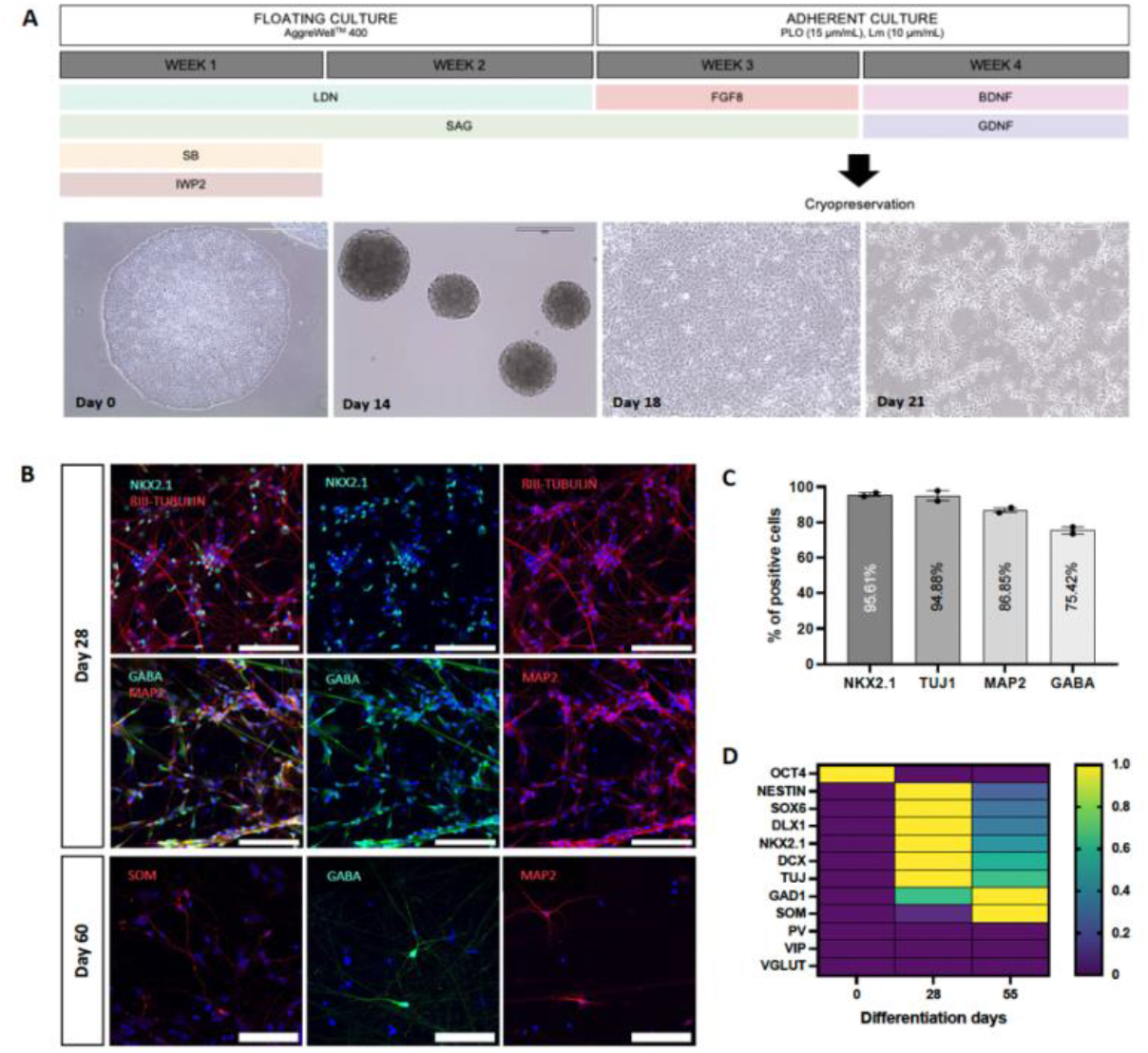
MGE-derived IN progenitor differentiation. **A**. Schematic representation of the differentiation protocol and pictures corresponding to different timepoints along the four weeks of differentiation. During the first two weeks, cells were maintained as floating spheres for neuronal induction and ventralization. At day 14 of differentiation, cells were plated and further differentiated into MGE-progenitors as adherent cultures. **B**. Representative IF images of IN progenitor markers after 28 days of differentiation. **C**. Quantification of the percentage of cells labeled by IN progenitor markers at day 28. **D**. Representative images of mature IN markers in 60-day old cultures. E. Transcript expression of key developmental genes in iPSC (day 0), IN progenitors (D28) and INs (D55). Scale bar = 400 μm (Day 0), 200 μm (Day 14, 18 and 21) and 100 μm (IF images).

For validation of *CHRFAM7A* null genotype in the WTSli002-A cell line, long amplification of the *CHRFAM7A* intron (exon A to exon 5) was carried out with the LongAmp® Taq DNA Polymerase Kit (ref. E5200S, New England Biolabs), according to manufacturer’s instructions. The PCR product was run in a 1% agarose gel to determine presence / absence and amplicon size.

### 2.8. Immunofluorescence (IF)

For immunofluorescent studies, cells were washed twice with PBS 1X and fixed with 4% paraformaldehyde (PFA) for 10 min at RT. After one washing in PBS1X, cells were incubated in a PBS 1X solution containing 0.1% Triton X-100 and 10% horse serum at room temperature (RT) for 10 min to block nonspecific binding. The primary antibodies, diluted in blocking solution containing 2% horse serum in PBS1X, were added overnight at 4°C. Three washings in PBS1X were performed and the secondary antibodies diluted in blocking solution were incubated for 2h at RT. An additional incubation with DAPI diluted in PBS1X was performed for 10 min at room temperature. The list of antibodies and dilutions used for IF experiments are presented in Suppl.Table S3.

### 2.9. α-bungarotoxin (α-BGT) binding

INs were washed twice with PBS 1X and fixed with 2% PFA for 15 min at RT. PFA was removed and wells were washed once with PBS 1X. To block unspecific binding, the cells were incubated with a PBS 1X solution containing 5% Horse Serum and 1% gelatin (ref. G7765, Sigma) for 1h at RT. To achieve a 1μg/mL concentration (∼125 nM), fluorescent α-BGT (ref. B35450, Invitrogen) was diluted 1:1000 in a PBS 1X solution containing 1% Horse Serum and 0.2% gelatin and incubated for 2h at RT. At this concentration, α-BGT is expected to saturate all α7-nAChR ligand binding sites at the cell surface. To control for α7-nAChR specific binding, 1 mM nicotine was added to the solution containing the toxin (non-specific binding, NS). The wells were rinsed two times and an additional incubation with DAPI diluted in PBS 1X was performed for 10 min at room temperature.

For quantification of α-BGT binding, the background intensity of each image was subtracted from the total intensity of fluorescence and divided by the number of cells. The number of cells per image was automatically counted with the StarDist plugin in ImageJ (Schmidt et al., 2018).

### 2.10. ROS Detection

For detection of ROS, cells were plated in Bio-One CELLSTAR μClearTM 96-well plates (ref. 655090, Greiner). The day of the experiment, 1 μL of CellRox deep red reagent (ref. C10422, ThermoFisher) was dissolved in 90 μL of B27GB medium to get a 25 μM stock. 2 μL of diluted CellRox reagent were added to one well containing 100 μL at a final concentration of 500 nM. The plate was protected from light and incubated for 30 min at 37°C. Following, the 96-wells were washed twice with warm DPBS and 100 μL of live Imaging solution (ref. A14291DJ, Invitrogen) was added per well to decrease background originated from phenol red-containing medium. 30 fluorescence reads at an interval of 1 min were acquired with a microplate reader (TECAN Spark, Chemogenomic and Biological Screening Platform, Institut Pasteur). The 6 consecutive cycles with more stable fluorescence were averaged and used for analysis. For each experiment, all fluorescence values were normalized to the condition containing α7-INs without Aβ treatment (Aβ0).

### 2.11. Imaging

Images were acquired with a confocal laser-scanning microscope (Zeiss LSM 700, Photonic BioImaging platform, Institut Pasteur) using a 20x or 40x oil objective, and with 488 nm and 647 nm laser for fluorescence excitation. To cover cells and their projections, Z-stacks were acquired.

### 2.12. Statistical analysis

Statistical analyses were performed using GraphPad Prism Version 9 software (GraphPad, San Diego, CA, USA) and are reported in each figure legend. Confidence levels of 95% were used.

## 3/ Results

### 3.1. iPSCs differentiate into MGE-derived IN progenitors in-vitro

A fully sequenced iPSCs cell line (WTSlii2, Wellcome trust) was used for these experiments. We tested and adapted a protocol to differentiate iPSCs into MGE-derived IN progenitors including a convenient step of cryopreservation at the end of the third week (Fig.1A). After four weeks of differentiation the majority of cells presented an IN progenitor phenotype (Fig.1B). 95.6% of the cells expressed the IN progenitor marker NKX2.1, 95% expressed the young neuronal marker Tuj1, 87% were positive for the neuronal marker MAP2 and 75% of the cells expressed GABA (Fig.1C). After 60 days in culture, mature IN markers like Somatostatin were detected, together with GABA and MAP2 (Fig.1D). Other mature IN markers such as PV and VIP were not detected under our experimental conditions.

Further characterization of the developing progenitors’ transcriptomic profile is shown in Fig.1E. While the pluripotency marker OCT4 was expressed in the iPSCs, the expression level was downregulated in 28- and 55-day old INs. As predicted, young neuronal markers (Nestin and Tuj1) and IN-specific progenitor markers (SOX6, DLX1, NKX2.1 and DCX) were mostly expressed in 28-day old INs, downregulated in 55-day old INs and not expressed in iPSCs. More mature markers of inhibitory IN phenotype like GAD1 and SOM were mostly upregulated in 55-day old INs and to a lesser extent, in 28-day old INs. Finally, other mature markers like PV, VIP and the glutamatergic marker vGLUT1 were not expressed at any of the timepoints studied (Fig.1E).

### 3.2. CHRFAM7A overexpression induces a reduction of α7-nAChR binding sites at the cellular membrane and a reduction of NACHO chaperone expression

The number of *CHRFAM7A* copies was tested in 4 different iPSC lines (WTSlii2, WTSlii8, WTSlii71, WTSlii30, Wellcome Trust) identifying a line with a null genotype (WTSlii2) (Suppl. Fig.1). To study the role of dupα7 in an isogenic background, we used a lentivirus to transduce the WTSlii2 cell line with the human *CHRFAM7A* gene and the fluorescent protein TdTomato (Suppl. Fig.2A). We performed cell sorting and iPSC cloning to ensure consistent *CHRFAM7A* gene dosage in all cultured cells.

Following four weeks of differentiation, INs derived from dupα7 transduced iPSCs (α7-dupα7-INs) expressed the *CHRFAM7A* transcript (Fig.2B). Although the overexpression of the human gene did not affect mRNA expression levels of *CHRNA7*, we observed a significant reduction in α7-ligand binding sites at the cell membrane (α-BGT-binding sites) of α7-dupα7-INs, suggesting a post-translational modulation (Fig.2C). In accordance with previous studies, a reduction of α-BGT-binding sites can be explained by the retention of the immature α7 protein in the ER, or by a reduction of ligand binding sites in heteropentameric receptors formed by both α7 and dupα7 subunits. While the two hypotheses are possible and non-exclusive, we focused on exploring further the alterations in α7-nAChR trafficking.

**Figure 2.**
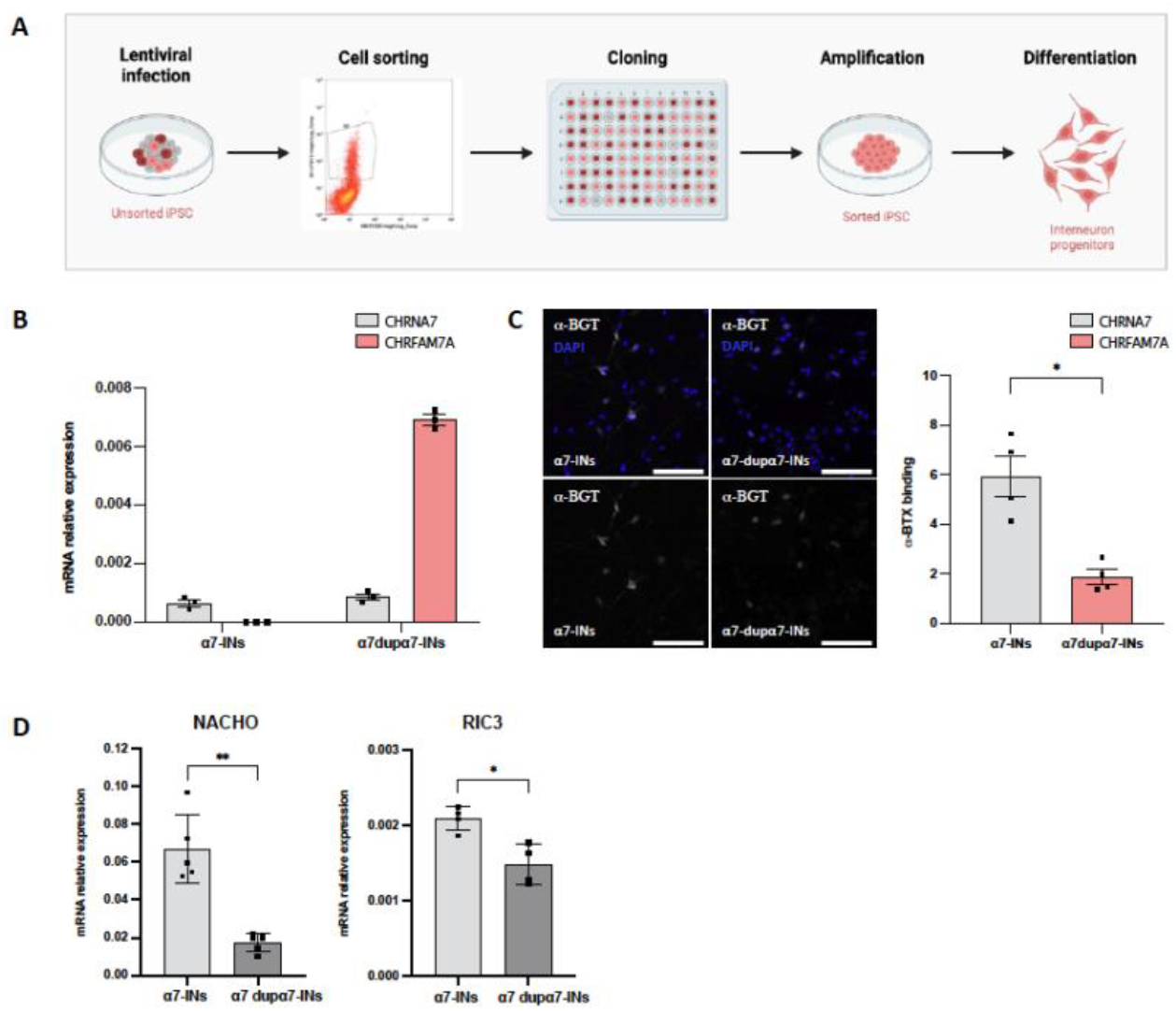
Overexpression of CHRFAM7A impairs α7-nAChR trafficking. **A**. Illustration representing the experimental steps from lentiviral infection to the obtention of INs containing equivalent levels of *CHRFAM7A* gene dosage. **B**. mRNA relative expression of *CHRFAM7A and CHRNA7* genes in α7-INs and α7-dupα7-INs. **C**. Representative images of cell surface α-BGX binding and quantification of total fluorescence. **D**. mRNA relative expression of the α7-nAChR chaperones NACHO and RIC3 in α7-INs and α7 dupα7-INs. mRNA levels are expressed as the fold change to the house keeping gene. α-BGT binding is expressed as adjusted fluorescence (total intensity – background intensity) divided by the number of cells. Scale bars = 100 μm. Data are presented in columns with mean values and SEM. For mRNA studies, each dot represents an independent culture and mRNA extraction. For α-BGT binding studies, each dot represents an independent experiment. For each independent experiment, the fluorescence of three images from different fields of view, is averaged. Statistical analyses were performed using Mann-Whitney test.

Two resident chaperones in the ER, NACHO and RIC3, play a critical role in assembling α7 subunits and facilitating their transport through the Golgi apparatus to the cell surface. While RIC3 has a moderate impact on α7nAChR trafficking in mammalian systems, NACHO’s involvement is essential to the process. Intriguingly, we found a significant reduction in the expression of both RIC3 and NACHO in α7-dupα7-INs when compared to INs not expressing the human subunit (α7-INs) (Fig.2D).

### 3.3. Nicotine treatment increases α-BGT-binding sites

Nicotine is a low-affinity α7-nAChR agonist that can bind to cell surface receptors but can also penetrate the cell membrane. Intracellularly, nicotine exhibits chaperone-like properties, as demonstrated by its ability to upregulate other nAChR subtypes, including α4ß2, which has a high affinity for the agonist.

In α7-IN cultures, exposure to nicotine for 30 min induced a slight but significant increase of cell surface α-BGT binding. However, these effects were less pronounced after 3 days of exposure, although a non-significant trend was still observed (Fig.3A, B). As previously observed, α7-dupα7-INs expressed significantly lower α-BGT-binding sites when compared to α7-INs (p-value: 0.0009, comparison not shown). Unexpectedly, nicotine exerted a highly significant upregulation of α-BGT-ligand binding sites in α7-dupα7-INs, mostly evident after 30 min of exposure but still conserved after 3 days of exposure to the agonist (Fig.3C, D). Interestingly, after acute nicotine treatment α7-dupα7-INs contained higher α-BGT-binding sites than α7-INs (p-value: 0.028; data not shown) but the levels were equivalent between the two cell cultures after longer nicotine exposure.

**Figure 3.**
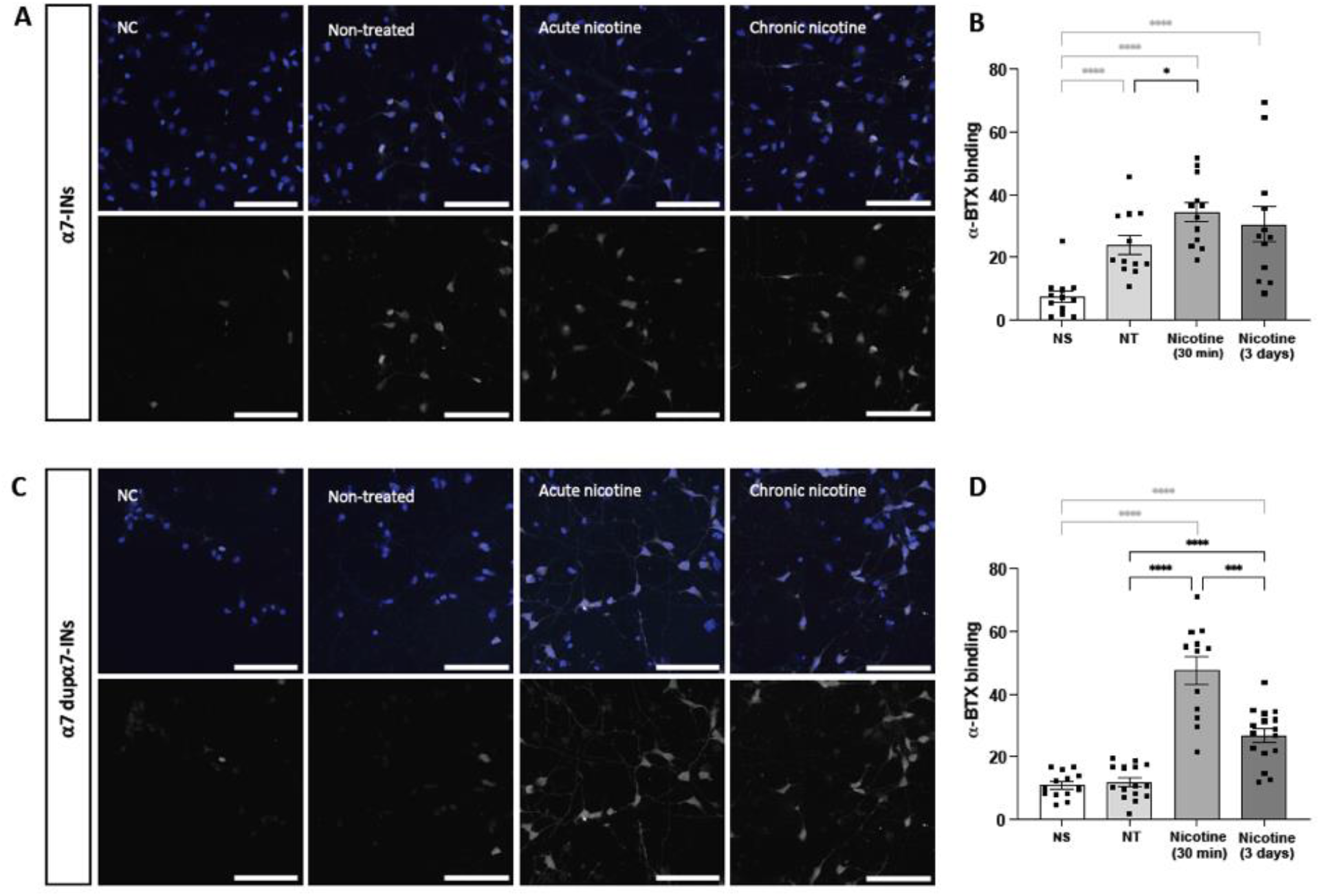
Nicotine treatment induces upregulation of surface α7-nAChR α-BGT binding sites. Representative images of fluorescent α-BGT binding under the four experimental conditions in α7-INs (**A**) and α7-dupα7-IN (**C**) cultures. Scale bars = 100 μm. B and **D**. Quantification of α-BGT binding expressed as adjusted fluorescence (total intensity – background intensity) divided by the number of cells, in α7-IN (**B**) and α7-dupα7-INs (**D**) cultures. Each dot represents a field of view of INs cultured in two different coverslips from one experiment. Non-specific binding (NS) refers to cultures not exposed to nicotine with post-fixation α-BGT binding displaced by nicotine. Non-treated (NT) condition refers to cultures not exposed to nicotine with post-fixation α-BGT binding. “Nicotine 30 min” and “3 days” refers to nicotine exposed cultures with post-fixation α-BGT binding. Statistical analyses were performed using Mann-Whitney test.

### 3.4. Differential response to Aβ_1-42_ peptide in α7-dupα7-INs and α7-INs

Previous studies have demonstrated that the Aβ_1-42_ peptide can induce toxic effects in neurons via α7-nAChR-dependent mechanisms (Szigeti et al., 2020; and reviewed in Ihnatovych et al., 2024). However, it has been suggested that at low doses, the peptide can activate the receptor and potentially play a neuroprotective role (Lasala et al., 2019). To investigate the impact of Aβ_1-42_ on cellular stress levels, we assessed the presence of reactive oxygen species (ROS) in the IN cultures after exposure to varying doses of oligomeric stabilized Aβ_1-42_.

Our results showed that α7-dupα7-INs had lower baseline levels of ROS compared to α7-INs. In our system, high doses of Aβ_1-42_ induced a slight but increasing trend of ROS in α7-INs but not in α7-dupα7-INs, suggesting a possible protective role of dupα7 in Aβ-induced oxidative stress response. Additionally, exposure to picomolar concentrations of Aβ_1-42_ significantly decreased ROS in α7-INs (p-value: 0.0079) to similar levels as those found in α7-dupα7-INs.

These findings support the idea that while low doses of Aβ_1-42_ may carry neuroprotective effects, higher doses may induce neuronal toxicity (Fig. 4A).

**Figure 4.**
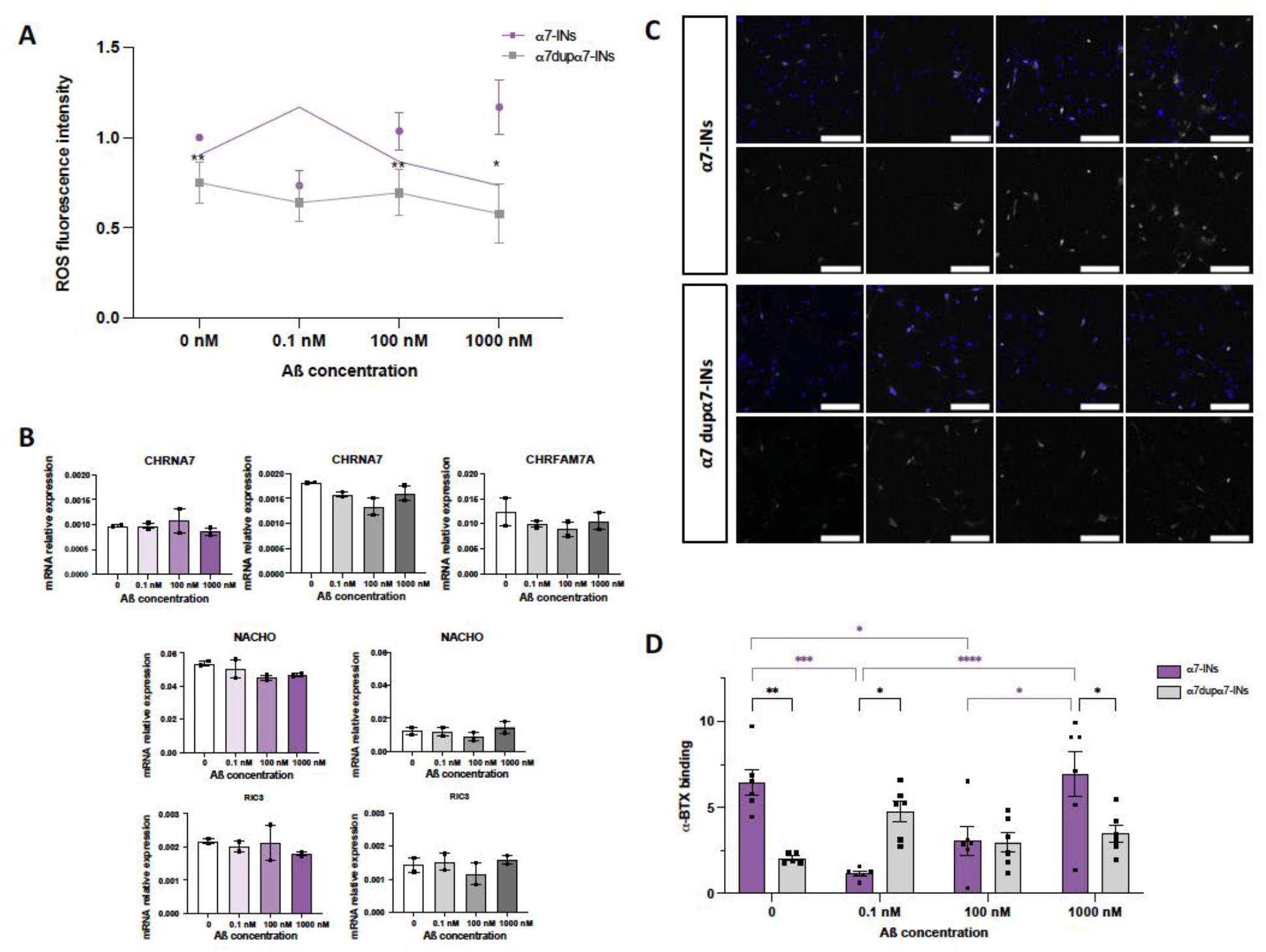
Differential effects of Aβ_1-42_ peptide on the production of reactive oxygen species, cell surface α7-nAChR ligand binding sites and transcript expression in α7-dupα7-INs and α7-IN cultures. **A**. Quantification of ROS-fluorescence intensity. Data is presented as the average and SEM of four independent experiments, normalized to the α7-INs control condition. Statistical analysis was performed using Mann-Whitney test. **B**. mRNA relative expression of *CHRNA7, CHRFAM7A, NACHO and RIC3* genes in α7-INs (purple scale) and α7-dupα7-INs (grey scale). **C**. Representative images of fluorescent α-BGT binding after 2h incubation of Aβ_1-42_ peptides. Scale bars = 100 μm. **D**. Quantification of α-BGT binding expressed as adjusted fluorescence (total intensity – background intensity) divided by the number of cells. Each point represents a field of view of two independent experiments. Data points were tested for normality (Shapiro-Wilk) and homoscedasticity (Levene’s test) and comparisons between groups were carried out with two-way ANOVA.

To investigate the potential mechanisms underlying the protective effects of dupα7, we examined whether Aβ_1-42_ peptide could regulate the receptor’s expression levels. However, we did not observe any differences in the transcript levels of α7-nAChR or its chaperone in response to Aβ_1-42_ exposure in either α7-INs or α7-dupα7-INs (Fig. 4B). We then explored potential post-translational mechanisms and found that the levels of surface α-BGT-binding sites in α7-dupα7-INs did not significantly change in response to Aβ_1-42_ exposure. In contrast, picomolar doses of Aβ_1-42_, and to a lesser extent, 100 nM doses, induced a significant decrease in surface α-BGT-binding sites in α7-INs (Fig. 4C, D). These results suggest a possible mechanism by which Aβ_1-42_ regulates accessible α-BGT-binding sites, with low doses reducing the available sites on the cell surface.

Further experiments are needed to elucidate the underlying cellular pathways responsible for these processes.

## 4/ Discussion

In this study, we examined the impact of the *CHRFAM7A* gene product on the regulation of α7-nAChR expression and its response to nicotine and Aβ_1-42_. To achieve this, we focused on investigating the role of *CHRFAM7A* in a controlled isogenic background. We identified a hiPSC line with a null genotype for *CHRFAM7A* and used lentiviral vectors to induce its overexpression. Additionally, we differentiated the hiPSCs into MGE-derived INs, which served as a relevant cell type for studying the effects of Aβ_1-42_. Similar to previous reported studies, the overexpression of *CHRFAM7A* led to a reduction in the cell surface α7-nAChR binding sites for α-BGT. Such reduction has been previously attributed to the formation of α7-dupα7 heteropentamers lacking binding sites for α-BGT and also to the retention of the α7 subunits at the ER, both leading to a decrease of expression of surface α7-nAChR (Araud et al., 2011; Liu et al., 2021; Lucas-Cerrillo et al., 2011; Maldifassi et al., 2018).

Assembly and trafficking of nAChRs are complex processes due to the diversity of nAChRs and factors involved including chaperone proteins (Colombo et al., 2013). More particularly, α7-nAChRs assembly within the ER and their trafficking through the secretory pathway are facilitated by the two chaperones RIC3 and NACHO (Castillo et al., 2005; Gu et al., 2016). In our experiments, we observed a significant downregulation of RIC3 and NACHO transcripts in α7-dupα7-INs. These data suggest possible defects in the regulation of α7-nAChR assembly within the ER and of their trafficking. These defects could be linked to the observed reduction of α-BGT binding sites in the cultured cells. It has been reported that the activity of NACHO requires the presence of three N-glycosylation sites of α7-nAChR (Kweon et al., 2020). As the dupα7 subunit lacks two of the three putative N-glycosylation sites, we could expect some impairments in the function of the chaperone NACHO in the cultured α7-dupα7-INs. RIC3 activity depends on its expression levels and neuronal cell compartments, dendrites vs. axons (Alexander et al., 2010). A dysregulation of α7-nAChR chaperones could be a cause or a consequence of the dysfunctional processing of α7-nAChR in the ER, leading to the observed reduction of α-BGT-ligand binding sites in the α7-dup α7-INs. It is important to note that mRNA levels do not always correlate with protein levels, and further experiments should be performed to confirm our data. Overall, these results support the idea of ER retention of the α7 subunit in α7-dupα7-INs.

It is well-known that nicotine can induce the up-regulation of nAChR receptors, as observed both in in-vitro experiments and in post-mortem analysis of brains of smokers (Barrantes et al., 1995; Benwell et al., 1988; Breese et al., 1997, Breese et al., 2000; Peng et al., 1997). This up-regulation is thought to occur through various mechanisms, including the mobilization of immature receptor pools within the ER (Corringer et al., 2006; Sallette et al., 2005). Although this mechanism has been mainly studied in high-affinity α4β2 nAChRs, α7-nAChR can also be up-regulated to a lesser extent (Barrantes et al., 1995; Breese et al., 2000; Peng et al., 1997). In agreement with these observations, we observed a slight but significant increase in α-BGT-binding sites in α7-nAChR, after 30 minutes of nicotine exposure. Interestingly, in α7-dup α7-INs, the amplitude of nicotine effect was stronger in α7-dup α7-INs as compared to α7-nAChR (Fig.3). This finding suggests that the presence of duplicated gene copies of the α7 subunit may modulate the response to nicotine exposure. The observed increase in α-BGT-ligand binding sites further supports the notion that nicotine could mobilize the intracellular pool of α7-nAChR and affect its availability at the cell surface. However, other mechanisms such as changes in receptor membrane stabilization or receptor conformational changes leading to variations in the α-BGT affinity cannot be excluded. To our knowledge, the nicotine-induced upregulation of α-BGT-binding sites in human cells expressing dupα7 has not been described before and could have significant implications in the context of the smoker population. However, the doses of nicotine used in our experiments are considerably higher than those typically encountered in smokers’ brains, and chronic exposure conditions may not accurately reflect the conditions found in nicotine consumers. It is therefore crucial to consider those aspects when extrapolating findings to in-vivo conditions. Interestingly, individuals carrying 2-3 copies of the *CHRFAM7A* gene have demonstrated a higher success rate in smoking cessation under treatment with the agonist varenicline of α7-nAChR, compared to those with only 0-1 copy of the gene. The variation in agonist-induced upregulation of α7-nAChR among individuals expressing different levels of dupα7 could potentially influence nicotine dependence and smoking cessation outcomes. Notably, while nicotine does not strongly activate α7-nAChR at concentrations typically found in smokers, varenicline activates the receptor at therapeutic concentrations in humans (Cameli et al., 2018).

Another aspect that we addressed in our experiments is the proposed protection of dupα7 subunit from Aβ induced toxicity (Ihnatovych et al., 2019). For this, we studied ROS levels in our system after exposure to three Aβ_1-42_ concentrations (picomolar, nanomolar and low millimolar). According to the suggested protective role of dupα7, we observed decreased levels of baseline ROS levels in α7-dupα7-INs and those were not affected by increasing Aβ_1-42_ concentrations. After 2h of exposure, α7-INs showed a slight increase of ROS production at higher doses of Aβ_1-42_. We could also observe a “protective” effect of picomolar doses of Aβ in α7-INs as shown by a reduction of ROS levels in these cells. It has been shown that Aβ can bind to α7-nAChR and elicit opposite effects according to its concentration (Pettit et al., 2001; Puzzo et al., 2008; Tozaki et al., 2002). While picomolar doses act as agonists of the receptor by rapidly desensitizing it, nanomolar doses bring the receptor towards a resting state (Lasala et al., 2019). Under our culture conditions, where endogenous α7-nAChR ligands are not present, we can expect that Aβ_1-42_ at picomolar concentrations acts as an agonist. Interestingly, under “non-agonist” conditions, α7-dupα7-INs exhibit lower levels of ROS compared to α7-INs. However, in the “Aβ-agonist” condition, ROS levels in α7-INs decrease to similar levels observed in α7-dupα7-INs. Although further pharmacological experiments are needed to confirm this hypothesis, dupα7 overexpression could induce signaling cascades leading to decreased ROS levels, independently of α7-nAChR signaling. Further investigation on regulatory ROS pathways including the PI3K/Akt/Nrf2 signaling pathway might shed light into this hypothesis.

Lastly, we investigated the levels of α-BGT binding sites after Aβ_1-42_ exposure in the two groups of INs. Although the specific site of Aβ binding to α7-nAChR is still unknown, it is believed to overlap with the ligand binding sites as it competes for binding with α-BGT (Cecon et al., 2019). Intriguingly, low doses of Aβ induced a decrease of α-BGTbinding sites in α7-INs. This reduction was significantly less important at 100 nM concentration and no significant changes were observed 1000 nM Aβ concentration or any other tested conditions in α7-dupα7-INs. There are several possible explanations for the decrease in α-BGT binding: increased degradation/endocytosis of the receptor, decreased trafficking of the receptor to the membrane, or decreased affinity for α-BGT binding. Considering our findings, a decreased affinity for α-BGT binding, could align with the proposed model. At picomolar concentrations of Aβ_1-42_, α7-nAChR rapidly undergoes a configuration change to a desensitized state with high agonist affinity (Lasala et al., 2019). Since INs were fixed before α-BGT binding, the receptors could be “frozen” in this desensitized configuration, resulting in lower α-BGT affinity. At higher doses of Aβ_1-42_, the receptors would be in a resting state configuration with lower affinity for Aβ_1-42_ and higher affinity for α-BGT (Lasala et al., 2019), which could account for our observations. However, these explanations are speculative and require further validation by other appropriate experimental techniques.

## Acknowledgments

We acknowledge the help of Dr. E. Bacchelli for providing us with a control cell line and are thankful for the scientific advice. The authors are thankful to Florent Haiss for scientific support. This research was funded by grants from the French National Research Agency ANR (ANR-17-NEU3-0004, iPS&BRAIN), from ERA-NET NEURON (iPS&Brain), from the “Fondation Alzheimer”, “Fondation Vaincre Alzheimer”, from “FRM Equipe 2019”, Institut Pasteur internal program “Projet Explore 2021”, and from ADPS-Allianz (to U.M., Head of Unit). This research was also funded by the Fondation pour la Recherche sur Alzheimer (to I.C-T.; C.T.’s PhD funding). The Pasteur-Paris University (PPU) international doctoral program provided the funding for M.L.P.’s PhD.

