## Supplemental Material for "*CHRFAM7A* overexpression in human iPSC-derived Interneurons dysregulates α7- nAChR surface expression and alters response to oligomeric β-amyloid peptide"

**Figure S1**

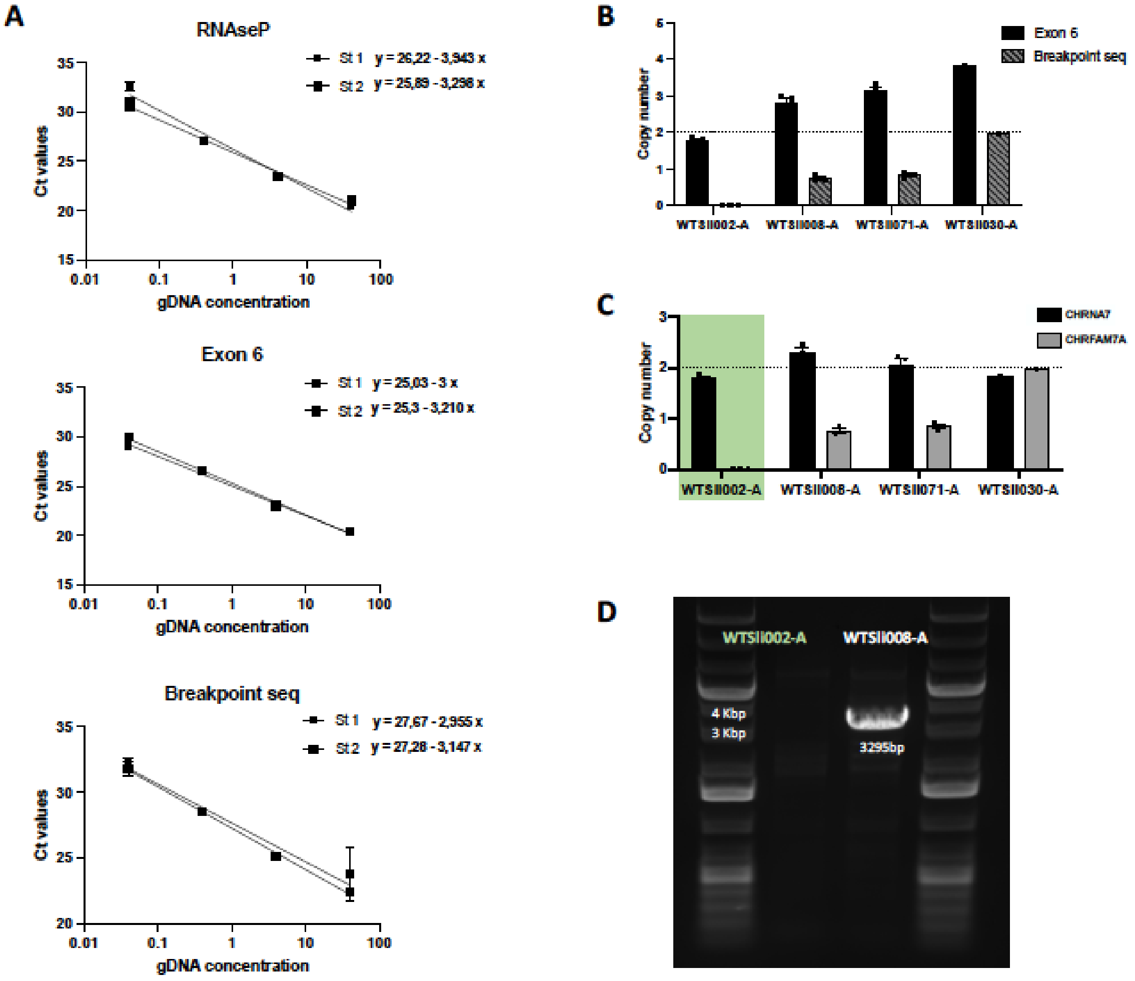


***Legend to Figure S1. CHRFAM7A CNV detection.*** **A.** Standard curves of Log10 gDNA dilutions and linear regression for primer efficiency calculation. **B.** Calculation of copy numbers of Exon 6 (present in both CHRFAM7A and CHRNA7) and breakpoint sequence (only present in CHRFAM7A) in four different iPSC lines. Ct values are compared to the control gene RNAseP (two copies) and to a control cell line (two copies). To calculate CNVs, a fold change of 1 to RNAseP is considered as 2 copies. **C.** CHRFAM7A and CHRNA7 copy number extrapolated from B. **D.** Amplification of CHRFAM7A specific intron (between exon A and exon 5) to validate null genotype of WTS2Ii002- A cell line.

**Figure S2**

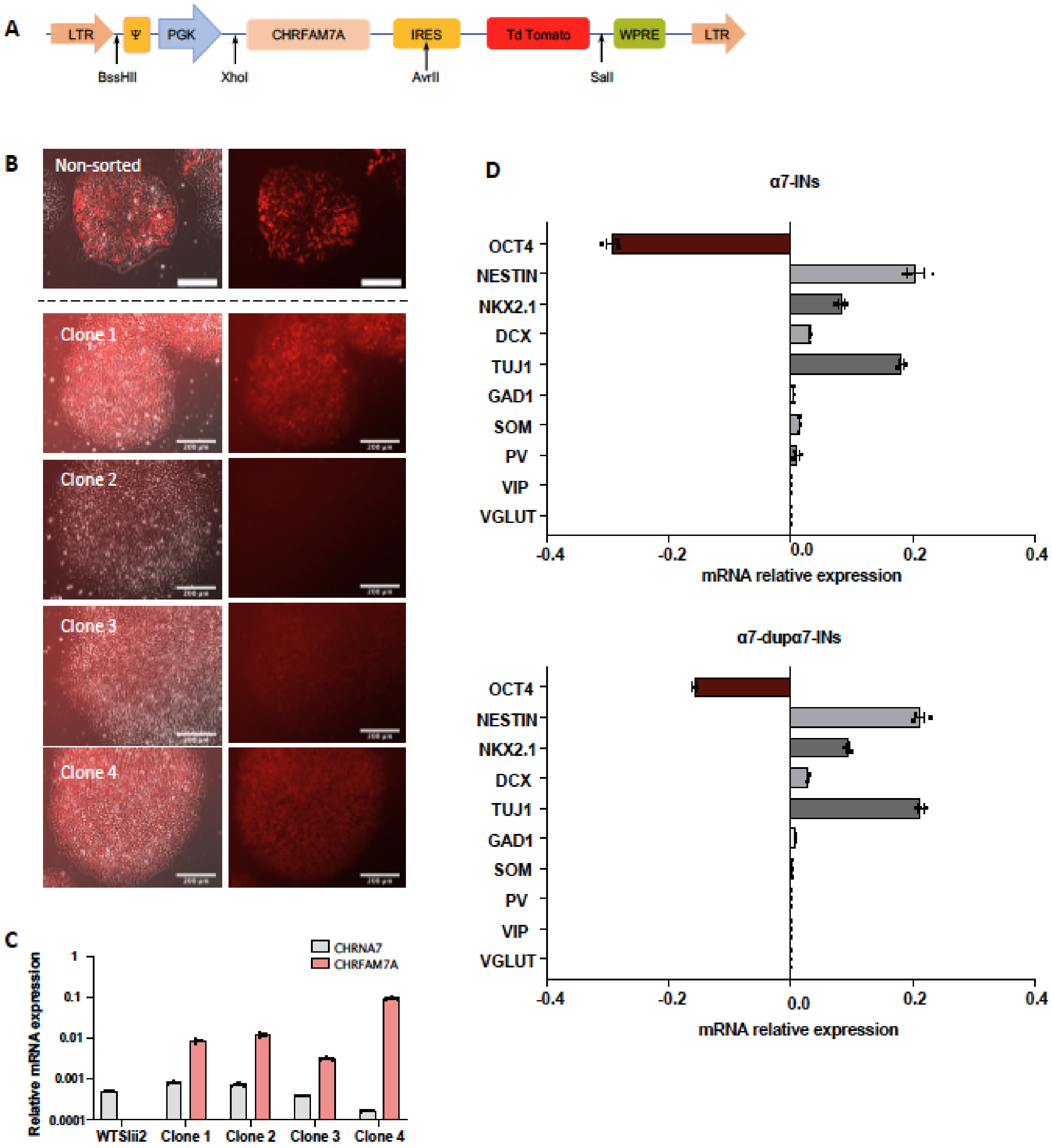


***Legend to Figure S2*. Lentiviral transduction and clone selection. A.** Lentiviral construct carrying *CHRFAM7A* and *TdTomato*. **B**. Representative images of iPSC colonies transduced with the lentiviral construct before cell sorting and after cloning. **C**. *CHRFAM7A* and *CHRNA7* expression in different iPSC clones expressed as fold change to the house keeping gene GADPH. The clone 3 was selected for the experiments presented in this article. **D**. Expression of IN differentiation markers in 28-day old INs, derived from control iPSC (top) and clone 3 (bottom). Similar expression patterns are shown in both control and transduced line, demonstrating no alteration of cell differentiation by the lentiviral construct.

**Supplementary Tables**

**Table S1:**

| Gene | Primer Forward 5' – 3' | Primer Reverse 3' – 5' |
|---|---|---|
| Oct4 | CAGCAGATCAGCCACATCGC | CCACACTCGGACCACATCCT |
| Nestin | TTCCCTCAGCTTTCAGGACCC | CTCAAGGGTAGCAGGCAAGG |
| Sox6 | ACAGCAAGAACAGATTGCGA | ATTGGGATCATGAGCGGAGG |
| Dlx1 | GGACTCACACAGACTCAGGTCAAGAT | GCCCTTCCCGGATGAAGAGTTA |
| Nkx2.1 | AAAGCACACGACTCCGTTCT | ATGAAGCGGGAGATGGCG |
| Dcx | AACTGGAGGAAGGGGAAAGC | CGGTCTCCAGTTTGATGGCT |
| Tuj1 | CTACAACGAGGCCTCTTCTCACAA | GGTTCCAGGTCCACCAGAATG |
| Gad1 | GCTAGAGATCTGCTTCCG | CAAATGTCTTGCGGACATAG |
| Som | CTTTAGGAGCGAGGTTCGGAG | ACTTGGCCAGTTCCTGCTTCC |
| Pv | GAGGAGGATGAGCTGGGATTC | CTTTAGGAGCGAGGTTCGGAG |
| Vip | GAAGCACCAGCAGGCAGTAA | CATTTCTGTGCCTCTCGCCC |
| vGlut1 | CGACAGCCTTTTGTGGTTCC | GGTTCATGAGTTTCGCGCTC |
| Chrna7 | CTGCAGATCATGGACGTGGATG | GAACTCTTGAATATGCCTGGAGG |
| Chrfam7a | CAAACTGCGATATTGCTGATGAGC | GAACTCTTGAATATGCCTGGAGG |
| Nacho | GCCTACAGTGAGATGAAACGTGC | CTGGTGGAAGAAGAGCACAGC |
| Ric3 | GAAATGCTATACTGCCATGCCTGG | CTTGCCTACGCTTGATACAGTCTG |
| APP | GGTAGAGTTTGTGTGTTGC | CTCTTCCTCTACCTCATCAC |

***Legend to Table S1*:** List of primers used for gene expression studies.

**Table S2:**

| Gene | Primer Forward 5' – 3' | Primer Reverse 3' – 5' |
|---|---|---|
| RNAseP | GCAAGTAAGTTTCTCCGAATCC | GCAAGTAAGTTTCTCCGAATCC |
| Exon 6* | GGGCATATTCAAGAGTTCCTGCTAC | CCACTAGGTCCCATTCTCCATTG |
| Chrfam7a breakpoint** | TCCTTGCCAATCAACTTTATGA | CACACCACCACACCTGGTTAAT |
| Chrfam7a Exon A-5 | GTGGATAGCTGCAAACTGC | GAA TGT GGC GTC AAA GCG |

***Legend to Table S2*:** List of primers used for CNV studies. *Exon 6 of both *CHRNA7* and *CHRFAM7A*. **Primer sequences from Cameli et al. 2018.

**Table S3:**

| PRIMARY ANTIBODIES | | | | |
|---|---|---|---|---|
| Name | Ref | Company | Host | Dilution |
| NKX2.1 | Ab133737 | Abcam | Rabbit | 1/2000 |
| ß3-TUBULIN | Ab9354 | Abcam | Chicken | 1/200 |
| GABA | A2052 | Invitrogen | Rabbit | 1/500 |
| MAP2 | MAB3418 | Millipore | Mouse | 1/200 |
| SOM | Sc55565 | Santa Cruz Biotechnology | Mouse | 1/250 |

| SECONDARY ANTIBODIES | | | | |
|---|---|---|---|---|
| Name | Ref | Company | Host | Dilution |
| **Anti-rabbit Alexa 488** | 711-545-152 | Jackson ImmunoResearch | Donkey | 1/500 |
| **Anti-chicken Alexa 647** | 703-005-155 | Jackson ImmunoResearch | Donkey | 1/500 |
| **Anti-mouse Alexa 647** | 715-605-151 | Jackson ImmunoResearch | Donkey | 1/500 |

***Legend to Table S3*:** List of antibodies used for immunochemistry.
